# SPARK: deciphering tumor-specific signaling networks through an integrative predictive model

**DOI:** 10.64898/2025.12.07.692871

**Authors:** Anson Zhang

**Affiliations:** Junior, BASIS Bilingual School Shenzhen, No.2 Xinsha Road, Shenzhen, China

**Keywords:** phosphoproteomics, oncogenic analysis, kinase-substrate associations (KSAs), machine learning (ML), protein language models, drug repurposing

## Abstract

While kinase-substrate associations (KSAs) are fundamental to cancer signaling, their rewiring patterns and functional roles across different cancers during cancer progression remain generally poorly characterized. Current predictive tools often fail to integrate three-dimensional structural constraints and cancer-specific dynamic information, limiting their biological relevance and translational potential. To address this, the present study constructs SPARK, a machine learning framework that integrates multi-modal features within cancer-specific contexts. These include data-driven features (such as phosphoproteomic co-expression, mutual information, protein-level abundance, and phosphosite-level abundance), knowledge-driven features (e.g., STRING protein-protein interactions), and structural features (ESM-2 embeddings compressed via an autoencoder). The training of the model was done with the adoption of the XGBoost classifier on data from eight CPTAC cancer cohorts. SPARK demonstrates strong predictive performance across different cancer types, with AUROC values ranging from 0.929 to 0.957 (mean: 0.944) in normal tissues and 0.941 to 0.958 (mean: 0.948) in tumor tissues. The accuracy of these predictions was further validated against experimentally determined kinase-substrate specificities. Using SPARK’s predictions, tissue-specific phosphorylation signaling networks are systematically reconstructed. These SPARK-derived networks reveal distinct phosphorylation patterns and kinase activation landscapes across cancers while identifying several highly tissue-specific kinases as promising therapeutic targets.

Graphical Abstract

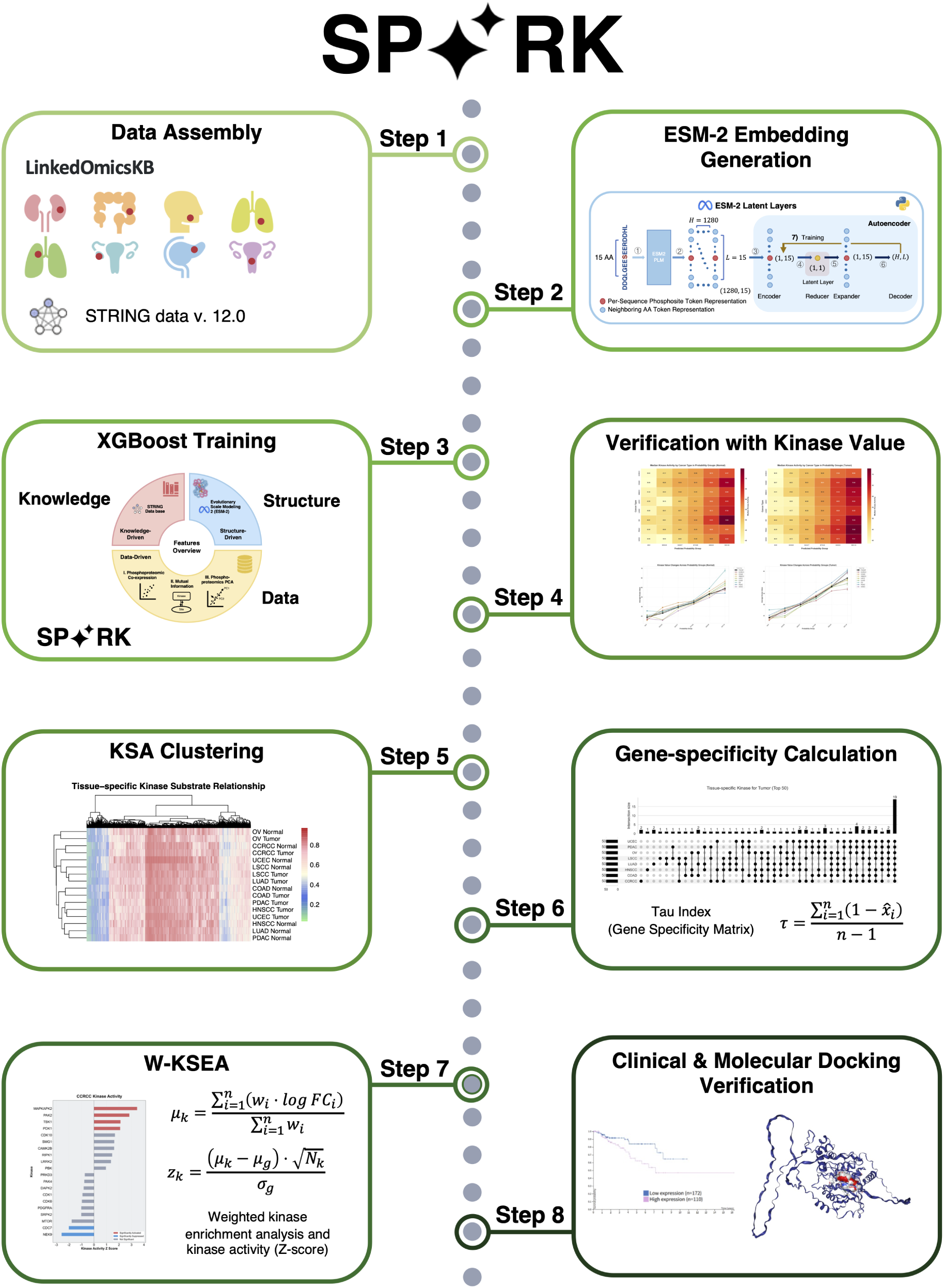

## 1 INTRODUCTION

Cancer progression is fundamentally a problem of signal rewiring in biological systems that allows the cell to bypass homeostatic controls on survival and growth [1]. Many of the effects of these changes map to protein phosphorylation, the most common and reversible post-translational modification (PTM) that control major cellular processes [2]. Dysregulated phosphorylation and aberrant kinase activity underlies hallmark cancer phenotypes, such as continuous cell division, resistance to apoptosis, and angiogenesis [3]. Accordingly, kinases have emerged as promising drug targets for cancer treatment [4]. As of May 2025, there are 88 FDA-approved small molecule protein kinase inhibitors since the clinical approval of imatinib in 2001 [5].

Leveraging protein kinases as potential drug targets for cancer depends on the understanding of kinase-substrate associations (KSAs) and their dynamic changes across cancer progression. Yet, fewer than 5% of human phosphosites are experimentally mapped to an upstream kinase, leaving a vast number of under-characterized KSAs [6]. The introduction of high-throughput mass spectroscopy (MS) and peptide microarray techniques has enabled unprecedented large-scale, rapid, quantification opportunities [7]. For example, Lancaster et al. were able to map 30,000 unique human phosphorylation sites in under 30 minutes and 81,120 unique phosphorylation sites within 12-hours of measurement [8]. These generated quantitative phosphoproteomic profiling has permitted computational based inference of KSAs.

Current KSA inference techniques mostly rely on the use of data- and knowledge-driven features based on expression values across samples, phosphosite amino acid (AA) sequence motifs, and annotation databases. Motif based technique include PhosX, a data-driven kinase activity inference method that compares the position-specified scoring matrix (PSSM) score of each phosphosite sequence against kinases to determine a similarity score that characterizes their association [9]. KINAID, an orthology-based kinase that supports multiple organisms, also uses PSSM similarity scores [10]. Other more comprehensive methods utilize supervised machine learning on a ground-truth dataset, often from PhosphoSitePlus (PSP), to enhance their predictions. The recently published SELPHI 2.0 employs a traditional random forest classifier that uses sequence motifs based on PSSMs, protein co-expression, phosphosite functional features, and kinase taxonomy information as features [11]. More advanced models include CoPheeKSA, a prediction algorithm using Extreme Gradient Boosting (XGBoost) model that integrates embedding features from CoPheeMap (a co-regulation classifier for phosphosite pairs) and KMap, PSSMs, and dynamic cross-cohort kinase-substrate co-expressions to achieve better pancancer KSA prediction [6]. Despite recent progress, these KSA-centric models are limited to data- and knowledge-driven perspectives that may hinder biological interpretability, and do not offer comprehensive translational relevance to cancer.

Phosphorylation, at its most fundamental chemical level, is a reversible PTM in which a phosphoryl group (*PO^−^*) is transferred to a substrate by cleaving the *γ*-phosphate from adenosine triphosphate (ATP) [12]. Given this, the structural interaction between the kinase and phosphosite and its neighbor is critical to understanding and deciphering this relationship. Current studies on KSA inference utilize data- and knowledge-driven features, lacking higher-order structural-driven support in their predictions. This results in limited biological interpretability given the 3D interaction between the kinase and phosphosite, leaving 3D interactions an unexplored opportunity to enhance KSA predictions. Furthermore, with the different nature of each cancer cohort, the construction of singular pan-cancer networks may not offer the most insight into using specific KSAs as drug targets. The construction of multiple tissue-specific heterogeneous networks can offer previously unanalyzed differences in cancer cohorts and offer better drug target opportunities. Importantly, cancer is a dynamic process that rewires inner phosphorylation pathways [13]. Thus, the implementation of differential kinase interaction networks (DKINs) on cancer can reveal changes in phosphorylation pathways caused by possible mutations or abnormal kinase activity, unveiling more accurate potential drug targets [14], [15].

Motivated by these opportunities, we constructed dynamic, tissue-specific differential networks that illuminate potential phosphorylation drug targets through data-, knowledge-, and structural-driven features in a machine learning model called SPARK. SPARK integrates data-driven features, including dynamic phosphoproteomic co-expression scores to characterize potential linear dependencies, pairwise mutual information scores to characterize potential nonlinear relationships, and scaled phosphoproteomics principal component analysis (PCA) to capture overall expression of the protein in the sample. STRING combined score was incorporated into SPARK as a knowledge-driven feature. STRING combined score is derived from prior knowledge based on genomic context predictions, high-throughput la experiments, conserved co-expression, automated textmining, and previous knowledge in databases [16]. STRING captures information that may not be represented in other data- or structural-driven features. Most importantly, SPARK captures structural information of phosphosites through Evolutionary Scale Modeling 2 (ESM-2) embeddings compressed through an autoencoder model. SPARK achieved exceptional results for all cancer cohorts in terms of area under the receiving operator curve (AUROC), area under the precision-recall curve (AUPR), and brier calibration scores. We then validated our results with experimentally derived kinase scores using the Kinase Library, demonstrating excellent prediction capabilities. Using the tissue-specific kinase-substrate differential network of eight cancers, including Clear Cell Renal Cell Carcinoma (CCRCC), Colon Adenocarcinoma (COAD), Head and Neck Squamous Cell Carcinoma (HNSCC), Lung Adenocarcinoma (LUAD), Lung Squamous Cell Carcinoma (LSCC), Ovarian Serous Cystadenocarcinoma (OV), Pancreatic Ductal Adenocarcinoma (PDAC), and Uterine Corpus Endometrial Carcinoma (UCEC), generated by SPARK, we first performed hierarchical clustering on predicted KSAs across different cancer cohorts. The clusters grouped tumor and mostly matched normal specimens by tissue of origin, demonstrating the capability of SPARK in capturing tissue-specific KSAs. Based on the prediction probabilities given by SPARK, we provide a set of high confidence kinase substrate associations (hcKSAs). Based on these hcKSAs, we generated kinase-specific and substrate-specific maps for the pan-cancer tumor cohort, revealing distinct patterns of the conserved expression of tissue-specific kinases and phosphorylation sites across multiple cancer types. Finally, we performed Weighted Kinase-Substrate Enrichment Analysis (W-KSEA) based on our predicted probabilities. We successfully identified kinases with increased activity in multiple tissues, several of which were potential therapeutic targets associated with tumor prognosis. Our results display the usefulness of our network in biochemical applications and new kinase drug target discovery.

## 2 RESULTS

### 2.1 SPARK: systematic phosphosite and kinase relationship prediction

To systematically predict the relationships between uncharacterized phosphosites and kinases, we present SPARK. SPARK is a tissue-specific, machine-learning-enhanced network that estimates probability of kinase-substrate interactions by utilizing data-, knowledge-, and structural-driven features (Fig. 1). By capturing a holistic frame of the proteome, SPARK is capable of generating effective predictions of kinase-substrate associations (KSAs) for understudied regions of the dark proteome.

**Fig. 1:**
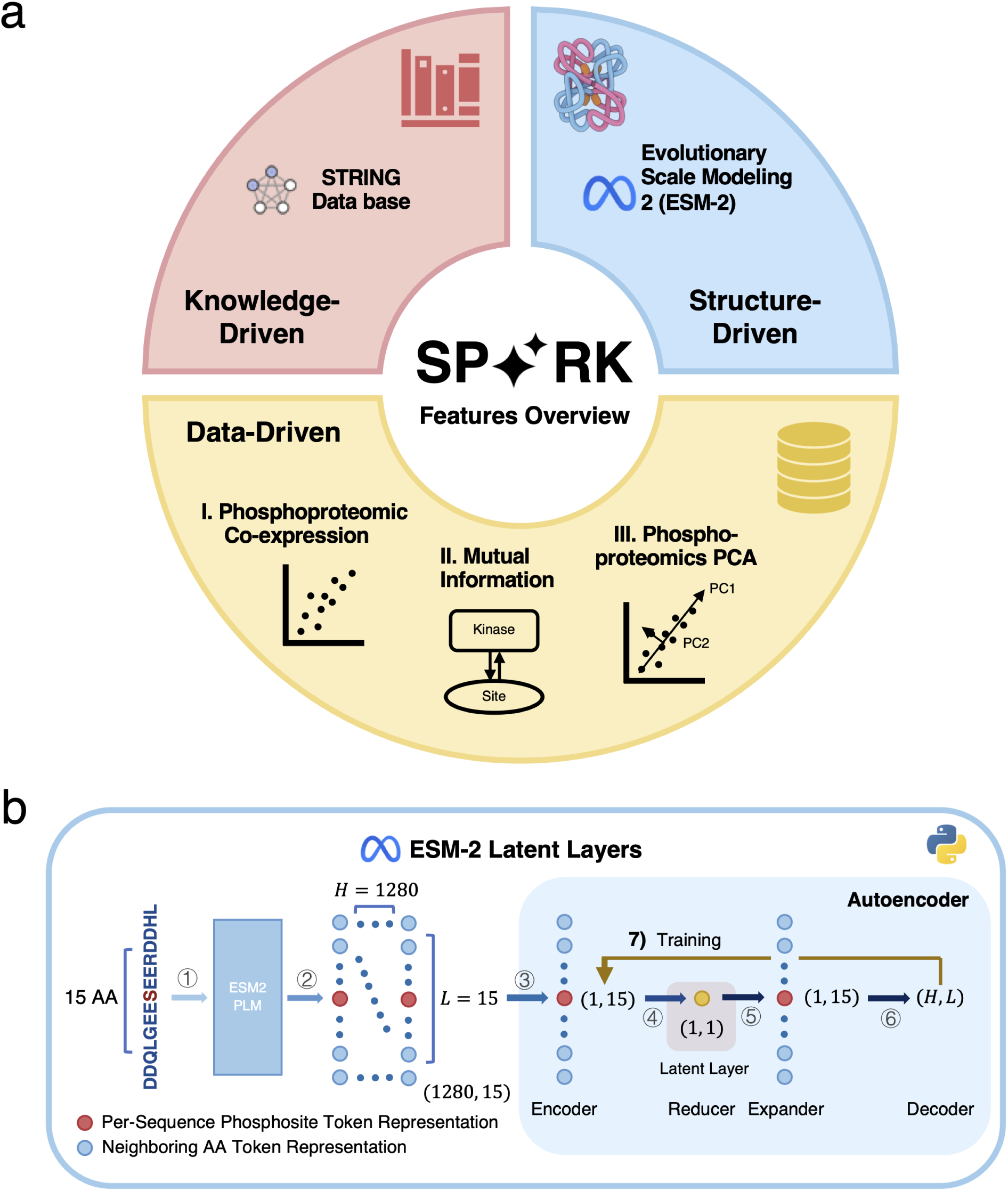
SPARK XGBoost classifiers and process. **a** Overview of the features used to train XGBoost. **b** Process used to obtain ESM-2 embeddings.

To develop the SPARK network, we utilized the Extreme Gradient Boosting (XGBoost) algorithm. Previously, XGBoost has proven extremely well-suited for this type of study. CoPheeMap, a pan-cancer co-regulation map of the human cancer phosphoproteome similar to that of SPARK, successfully employs XGBoost in KSA prediction. XGBoost offers several important advantages when compared to other traditional machine learning classifier algorithms. Firstly, XGBoost is capable of handling missing (NA) values, which is especially important in handling biological datasets where missing data is prevalent [6]. Secondly, XGBoost yields higher precision-recall performances, since it provides built-in mechanisms for coping with data imbalances [17]. Phosphoproteomic datasets are often severely imbalanced because known positive kinase-substrate pairs are rare compared to known negative kinase-substrate pairs [6]. Thirdly, compared to other traditional machine learning classifier algorithms, such as Random Forest, XGBoost has built in L1/L2 regularization, helping it prevent overfitting [17].

The tissue-specific SPARK XGBoost algorithm was constructed using three types of features: data-, knowledge-, and structural-driven features (Fig. 1a). Data-driven features integrate kinase-substrate pairwise phosphoproteomic co-expression, pairwise mutual information, and phosphoproteomic principal component analysis (PCA). These features create a holistic profile for each kinase-substrate pair based on expression values, representing the linear and nonlinear dependencies that may exist between them. Knowledge-driven features utilize STRING database combined scores. STRING combined scores provide prior-knowledge-based understanding of the kinase-substrate pair, given its incorporation of textmining and knowledge in previous databases [16].

To capture 3D structural features, we employed Evolutionary Scale Modeling 2 (ESM-2) embeddings, compressed using an autoencoder model. ESM-2 offers the advantages of 1) not requiring MSAs for capturing high resolution structural information and constraints 2) providing atomic-level projection accuracy in its structural predictions based directly from individual sequences [18]. The ESM-2 structural features were generated by extracting the latent layer of an autoencoder model. The autoencoder converged with a low mean square reconstruction error of 0.0126 and 0.0141 for tumor and normal ESM-2 embeddings, respectively. This illustrates the effectiveness of using autoencoders instead of other methods, such as mean pooling, in minimizing data loss and capturing a comprehensive structure of each phosphosite.

### 2.2 Model Performance and Probabilistic Calibration

To comprehensively evaluate SPARK’s ability to predict KSAs across diverse cancer types, we conducted a comprehensive, pan-cancer evaluation. The results demonstrate the excellent discriminative power and well-calibrated probabilistic prediction capabilities of SPARK. Across cancers, our XGBoost classifiers attained a high Area Under the Receiving Operator Curve (AUROC) ranging from 0.929 to 0.957 (mean: 0.944) for normal and 0.941 to 0.958 (mean: 0.948) for tumor (Fig. 2). The Area Under the Precision Recall Curve (AUPR) ranged from 0.797 to 0.896 (mean: 0.844) for normal and 0.834 to 0.881 (mean: 0.853) for tumor (Fig. 2).

**Fig. 2:**
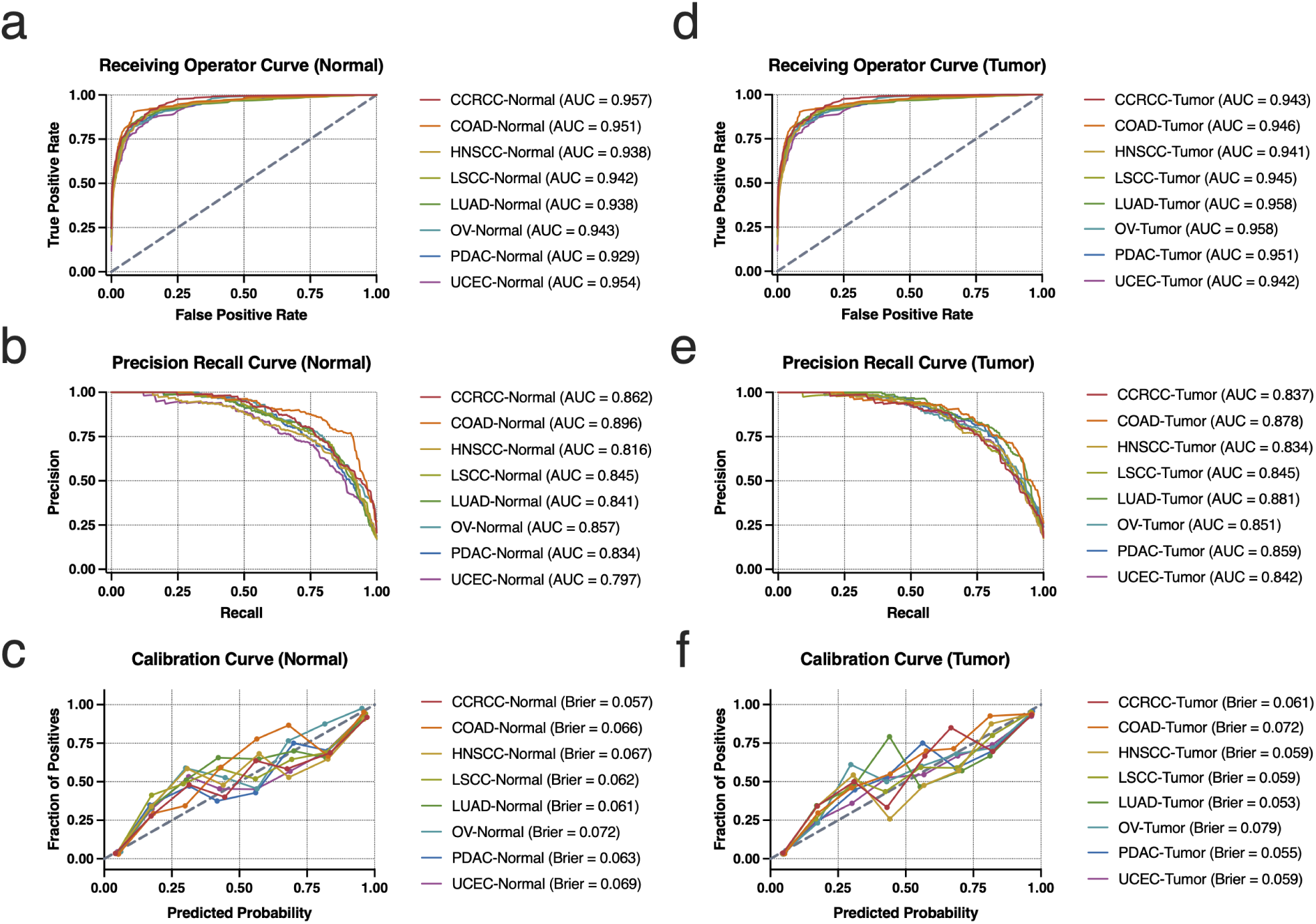
SPARK XGBoost classifier model performance and probabilistic calibration. **a-c** Normal cohort performance in terms of ROC (**a**), PR (**b**), and calibration curve (**c**). **d-f** Tumor cohort performance in terms of ROC (**d**), PR (**e**), and calibration curve (**f**).

A core advantage of our method is its output in probabilistic score, different to binary classifiers. To analyze the reliability of our probabilistic score, we analyzed the model’s probability calibration. As shown in Fig. 2c and Fig. 2f, SPARK demonstrates excellent calibration with low Brier scores (below 0.08 for all cohorts) and the calibration curve closely matching the diagonal, indicating high consistency between predicted probabilities and observed frequencies. This elevates our model beyond a conventional binary classifier, making it reliable in probabilistic estimates. High-confidence probabilistic estimates permit the ranking of KSAs likelihoods, which allows researchers to prioritize their research on follow-up experiments and offers them a solid basis for nuanced biological interpretation [19].

### 2.3 Probability-dependent prioritization of kinase-substrate relationships

To further validate the predictive efficacy and ranking capability of SPARK, we evaluated the percentile rank (kinase value) of predicted KSAs with the values from the Kinase Library. Our results demonstrate a significant positive association between our predicted probability and kinase values across cancer types in both normal (spearman correlation, *ρ* = 0.147, p-value *<* 2.22 × 10^-16^) and tumor (spearman correlation, *ρ* = 0.161, p-value *<* 2.22 × 10^-16^) cohorts. This verifies the reliability of SPARK in predicting KSAs. Furthermore, the heatmaps in Fig. 3a and Fig. 3b demonstrate prediction probabilities consistently corresponding to higher median kinase scores. This pattern holds true across diverse cancer contexts, highlighting the generalizability of the model’s ranking performance. Fig. 3c and Fig 3d. confirms the reliability of our model in prioritizing high-confidence KSAs. Notably, median kinase values in tumor cohorts appears generally larger than normal cohorts. This suggests that kinases in cancer cells are more active, which may reflect increased signaling associated with tumor progression [20].

**Fig. 3:**
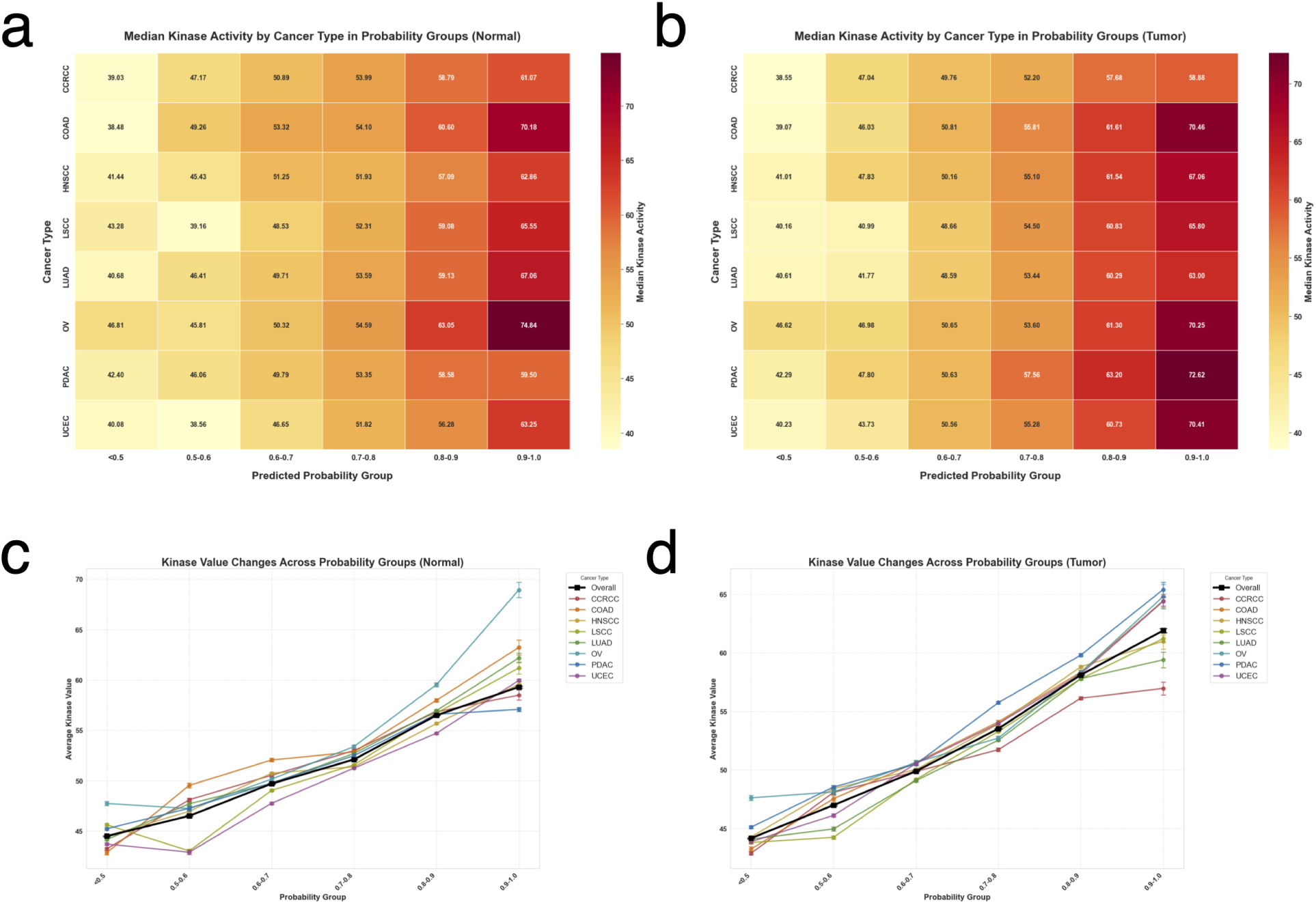
Relationship between kinase value and binned probability groups. **a-b** Median kinase value by tissue type in probability groups for the normal cohort (**a**), median kinase value by tissue type in probability groups for the tumor cohort (**b**). **c-d** Mean kinase value changes across probability groups with confidence intervals for the normal cohort (**c**), and mean kinase value changes across probability groups with confidence intervals for the tumor cohort (**d**).

### 2.4 Analysis of tissue-specific kinase-substrate associations in pan-cancer data

SPARK successfully delineates tissue-specific KSAs across multiple cancer types. We performed hierarchical clustering analysis on a randomly selected sample of 5000 kinase-substrate pairs. Tumor and normal groups were processed independently, and the results are shown in Fig. 4. As shown, groups were mostly clustered according to cancer type and not cancer state (tumor or normal). This is especially prominent in ovarian cancer (OV) and clear-cell renal-cell carcinoma (CCRCC). This exemplified tissue-specific characteristic is a fundamental principle in cell signaling, confirming that kinase activity largely depends on cancer type (https://www.nature.com/articles/ncomms1871).

**Fig. 4:**
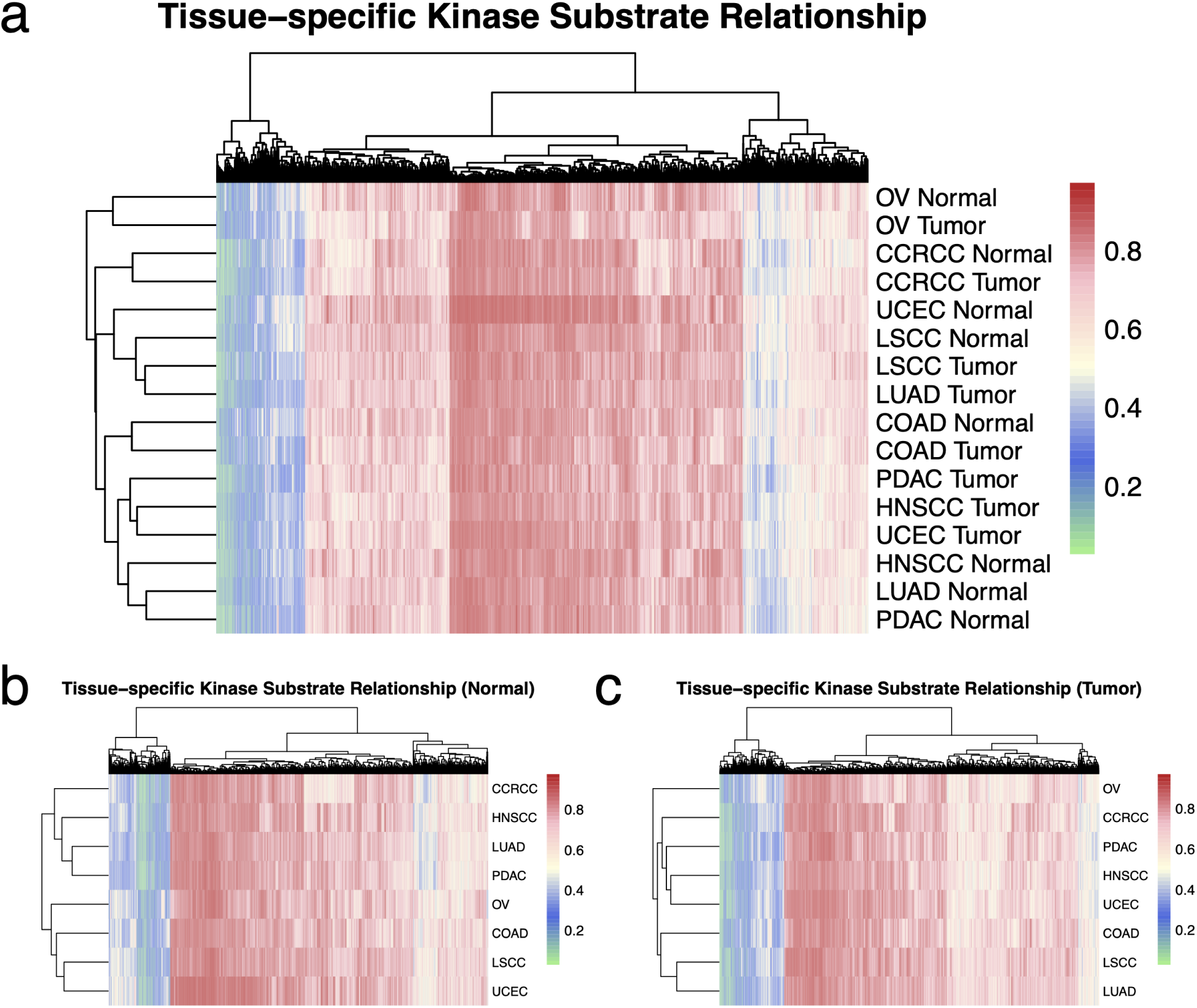
Relationship between kinase value and binned probability groups. **a** Tissue-specific kinase substrate relationship for pan-cancer data. **b** Tissue-specific kinase substrate relationship for normal cohorts. **c** Tissue-specific kinase substrate relationship for tumor cohorts.

### 2.5 Distinct patterns of kinase and substrate specificity across cancer types

To gain insight into the function of top priority kinases and phosphosites function in biological systems, we filtered out KSAS with the top 1% of probability and pair-wise kinase value *≥* 80% in the tumor cohort (see methodology). To understand kinase and substrate specificity, the tau (*τ*) index of each priority kinase and phosphosite was calculated based on count, and the results are visualized in Fig. 5. This analysis revealed fundamental differences between the specificity patterns of kinases and their target substrates.

**Fig. 5:**
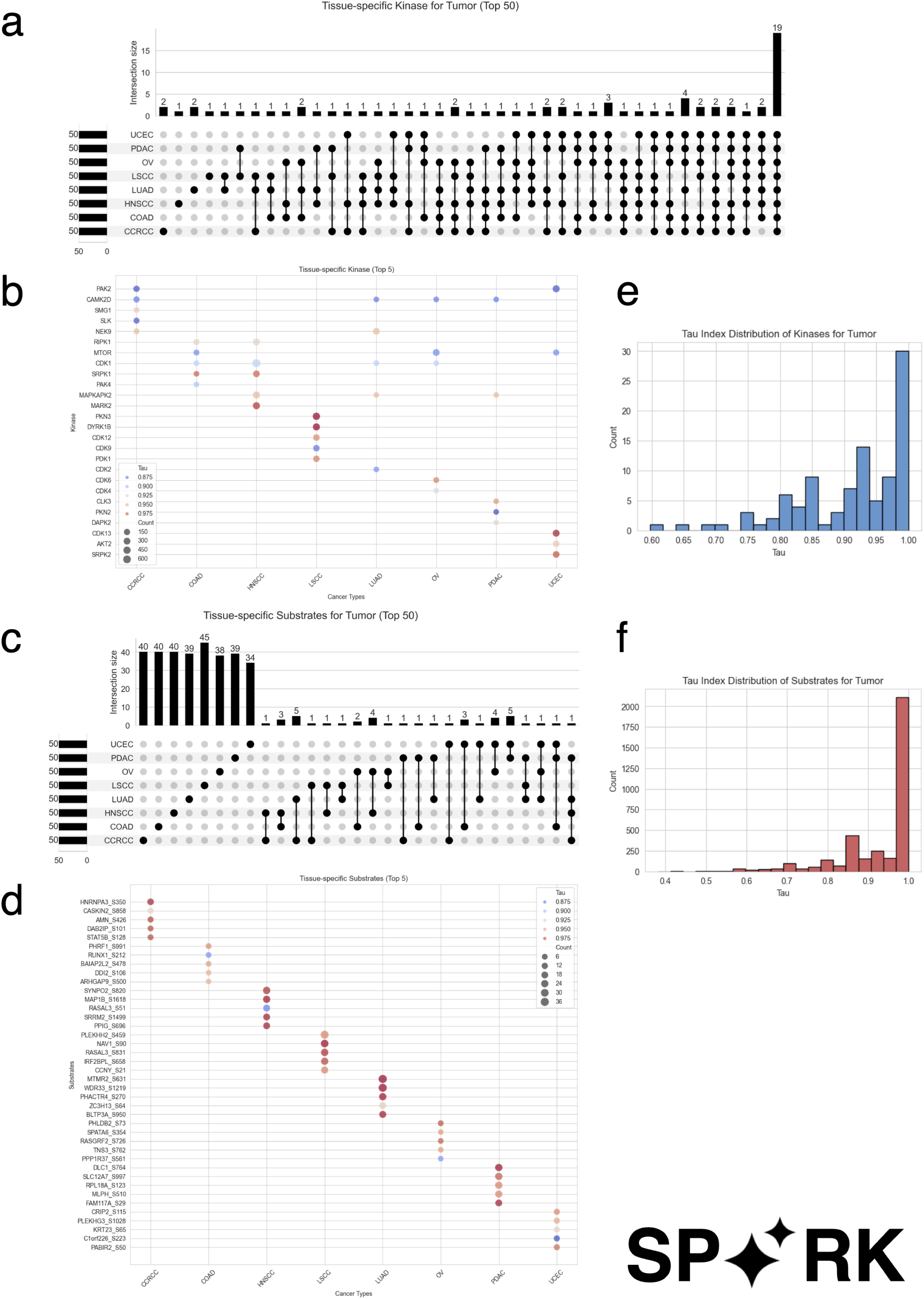
High specificity kinases and substrates across tissue types. **a, c** UpSet diagrams of top 50 tissue-specific kinases and substrates. **b, d** Top 5 tissue-specific kinases and substrates. **e, f** Tau index distribution of kinases and substrates.

The count and tau index of the top 50 kinases with the highest tau index across cancer types is shown in the upset diagram in Fig. 5a. As shown, the kinases demonstrate high conserved expression, with 19 core kinases exhibiting low specificity and appearing in all 8 tissues. This suggests the ubiquity of kinases across all cancer types and high functional importance. However, our kinase results also suggest some sparse difference in kinase specificity across cancer types. Specifically, LUAD, CCRCC, HNSCC, LSCC, all have some individual high specificity kinases (*≤* 2), suggesting the activation of some unique, tissue-specific, kinase-driven pathways. In contrast, analysis of the top 50 substrates in Fig. 5c reflects a pattern of low conserved expression across tissue types. This difference can be visualized in the histograms Fig. 5e and Fig. 5f, where substrates appear significantly more tissue-specific. This striking contrast suggests that while the core kinase group is largely shared across cancers, downstream signaling outputs are highly specialized and context-dependent.

To elucidate the difference in the patterns, we focused on the top 5 kinases and substrates in each tissue with the highest tau index (Fig. 5c and 5d). A comparison of Fig. 5c and 5d shows that even the most specific kinases are shared across tissue types, while the most specific substrates are entirely dominant to one tissue. For example, CAM2KD exhibits behavior in CCRCC, LUAD, OV, and PDAC.

Analyzing these distinct patterns allows us to select high priority kinase and substrate candidates that are highly specific, but further analysis on whether they exhibit enhanced functional activity in their respective tissue was still required. To this end, we performed weighted kinase-substrate enrichment analysis (W-KSEA).

### 2.6 Comprehensive analysis of kinase activity across cancers reveals PAK2 as key player in CCRCC and uniqueness of PDAC profile

Performing W-KSEA has allowed us to construct kinase activity profiles for the 8 tissues consisting of the top 20 kinases with the highest absolute value of kinase activity (Fig. 6. By systematically comparing the profile across the 8 cancer types, we revealed that most kinases with the highest kinase activity exhibited high conserved expression, since they acted as fundamental drivers of cancer itself, even though some kinases were highly tissue-specific. For instance, cell cycle-related kinases, including CDK1 and CDK2, were widely activated in most cancers with an average Z-score *>* 2.5, reflecting a common hallmark of cancer cell proliferation. CDK2 regulates entry into S phase (G1 checkpoint), while CDK1 regulates the start of mitosis (G2/M checkpoint), which are both key hallmarks of abberant activity shared in tumor cells [21]. Interestingly, the CDK family was significantly suppressed in Pancreatic Ductal Adenocarcinoma (PDAC), different to other tissues where CDK exhibit significant activation (Fig. 6f). CDK inhibition in PDAC is specifically related to vulnerability in KRAS addiction, a hallmark of PDAC [22]. Clear Cell Renal Cell Carcinoma (CCRCC) also exhbits an unqiue kinase activity profile, with its signaling network dominated by the p21-activated kinases (PAKs) and the MAPK pathway (Fig. 6a). PAKs have been clinically proven to be related to hypoxia, a hallmark of CCRCC [23].

**Fig. 6:**
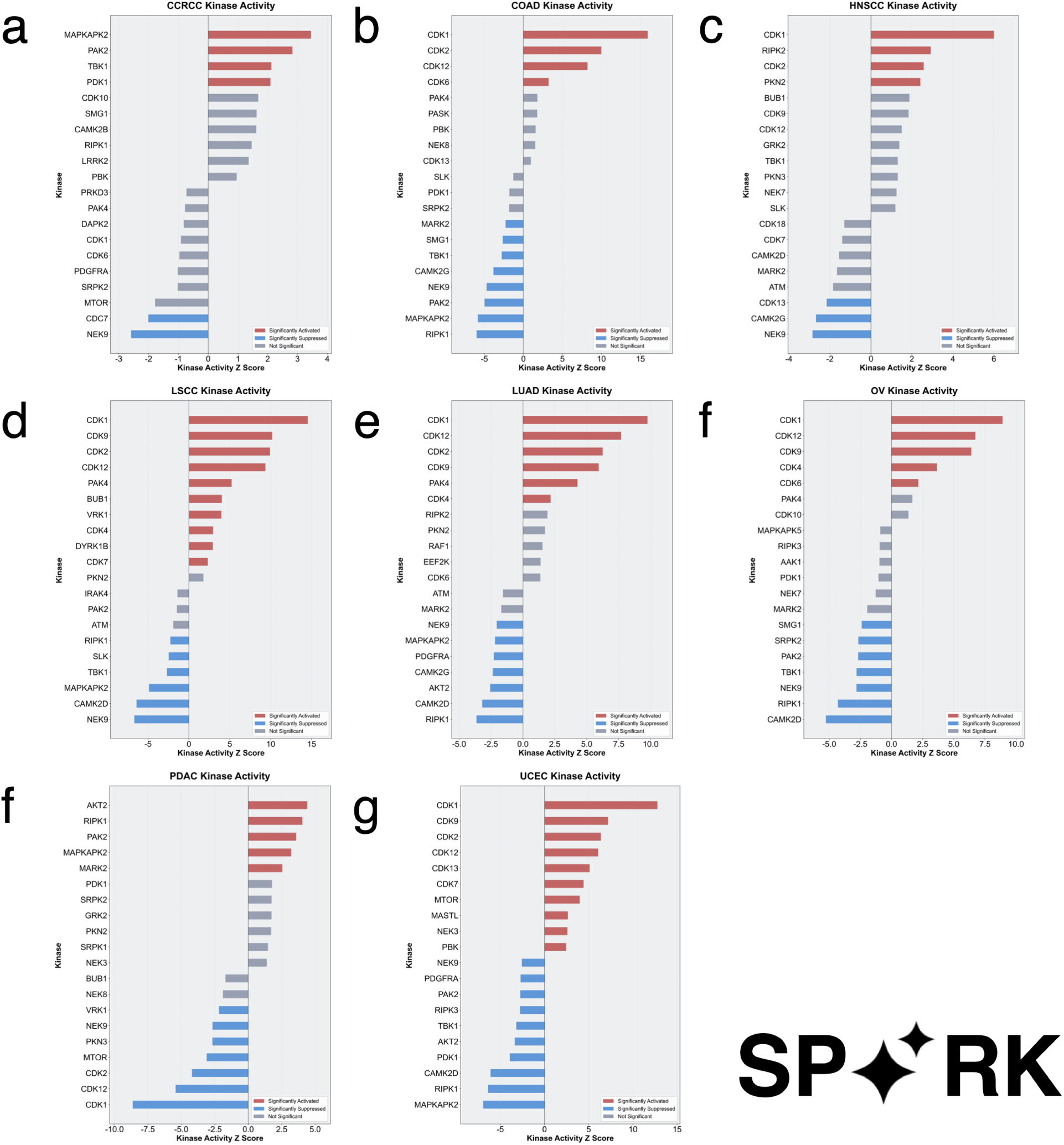
Top 20 kinases with the highest kinase activity. **a-g** Kinases for CCRCC, COAD, HNSCC, LSCC, LUAD, OV, PDAC, and UCEC.

In CCRCC samples, PAK2 exhibited the most significant increase in kinase activity (Z-score = 2.83, p-value = 4.6*×*10*^−^*^3^), with activation levels far higher than in other cancer types (such as Z-score = −4.97 in COAD). This specific activation is directly correlated with clinical outcomes. Data from the Kidney Renal Clear Cell Carcinoma obtained from the Human Protein Atlas showed that patients with high PAK2 expression had a 20% decreased 5-year survival rate (p-value = 1.2 *×* 10*^−^*^4^).

Given the central role of PAK2 in the pathogenesis and prognosis of CCRCC, we sought potential targeted therapeutic options. By querying the DrugBank database, we identified fostamatinib, an FDA-approved SYK inhibitor, as a potential PAK2 binder. Molecular docking simulations revealed a strong interaction between fostamatinib and the PAK2 kinase domain, with a Vinascore of −9.0 kcal/mol, indicating high binding affinity (Fig. 7b and Fig. 7c).

**Fig. 7:**
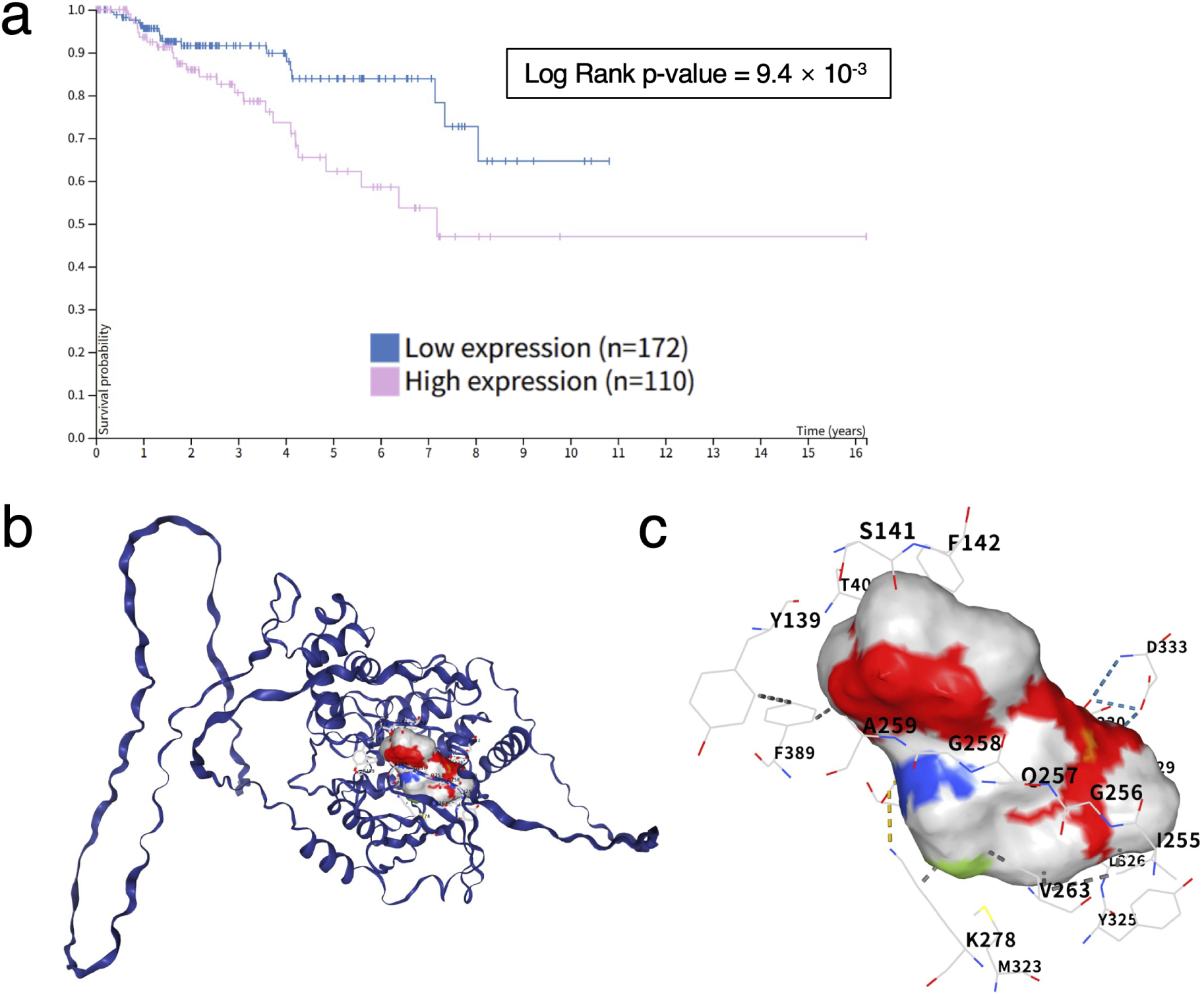
Key information regarding PAK2 in CCRCC. **a** Survival plot of PAK2 for Kidney Renal Clear Cell Carcinoma obtained from The Human Protein Atlas. **b-c** Molecular docking simulations between fostamatinib and the PAK2 kinase domain.

## 3 DISCUSSION

### 3.1 Advancing KSA prediction using SPARK

Our study introduces SPARK, the first systematic phosphosite and kinase relationship prediction framework that integrates data-, knowledge-, and structural-driven features. SPARK advances KSA prediction using 3D structural features represented via ESM-2 embeddings, dynamic phosphosite co-expression, STRING knowledge-based scores and tissue-specific signaling contexts. Existing ML based tools, such as SELPHI 2.0, CoPheeKSA, mostly rely on sequence motifs, kinase profiles, co-expression, and other data-driven features [6], [11]. SPARK uniquely and effectively leverages ESM-2 as a protein language model to capture structural information governing KSAs (Fig. 1). Furthermore, while common techniques in ESM-2 residue pooling employ mean pooling, SPARK utilizes a separately trained autoencoder for normal and tumor cases, attaining low reconstruction error and minimizing information loss during pooling (Fig. 1). The integration of structural representation enables SPARK to perform exceptionally well with unprecedented accuracy (mean AUROC 0.944 for normal and 0.948 for tumor) and probability calibration (Brier score *<* 0.08), critical for prioritizing high-confidence KSAs for experimental study and deciphering the “dark” proteome. We confirmed the high reliability of SPARK with experimental kinase activity derived from the Kinase Library, with significant positive correlation in both normal (spearman correlation, *ρ* = 0.147, p-value *<* 2.22 × 10^-16^) and tumor (spearman correlation, *ρ* = 0.161, p-value *<* 2.22 × 10^-16^) cohorts. We also confirmed the high confidence of our high-probability KSAs, demonstrated by the increase in median kinase value that corresponded with increasing probability groups (Fig. 3). Moreover, while most current models predicting KSAs produce a single pan-cancer heterogeneous network, SPARK is trained on each tissue and produces tissue-specific, differential, heterogeneous networks. This allows for more comprehensive review of potential KSAs of interest and cancer state (normal vs. tumor) comparison [14], [24].

### 3.2 Unveiling key information about tissue-specific KSAs in cancer progression with SPARK differential networks

Through hierarchical clustering of all tissues in a pan-cancer evaluation, we revealed that tissue groups were mostly clustered according to cancer type and not cancer state (normal vs. tumor). In the grip of this view, we see the importance of tissue type in determining cancer rewiring processes and kinase activation. However, the large difference in the clustering of certain tissues, such as uterine corpus endometrial carcinoma (UCEC), imply fundamental changes in the cell signaling pathway due to cancer that deviate significantly from the normal tissue [6]. We then analyzed the tissue-specificity of each priority kinase and substrate using the tau index. Interestingly, we found that phosphosites exhibits significant tissue-specificity when compared to kinases, which can be attributed to 1) phosphosites having numerical dominance over kinases 2) kinases being present and shared broadly across tissues, and phosphosites being more specific to the local signaling environment [1]. This underscores the importance of local contexts in downstream signaling outputs (substrates), under the influence of external factors such as accessory proteins. Furthermore, our oncogenic analysis of tissue-specificity also reveals a list of potential priority kinases and substrates that may be tissue-specific drivers of cancer, helping researchers prioritize their research in laboratories.

To further understand the functional importance of these tissue-specific priority kinases, we performed W-KSEA. W-KSEA is a newly developed technique based on kinase enrichment analysis that integrates the predicted probability of SPARK. This integration allows for the calculation of kinase activity scores on newly identified KSAs predicted by the model, allowing for its application to the dark proteome. This also reduces noise from low-probability edges and enhances biological relevance. We successfully identified CDK1 and CDK2, kinases that are related to the cell-cycle across all tissue types, demonstrating the biological validity of our SPARK [21]. We also revealed the uniqueness of the PDAC kinase profile with its significant inhibition of the CDK aligning with results in current studies [22]. Furthermore, we successfully identified p21-activated kinases (PAKs) as the unique kinase that dominates the CCRCC profile, which are linked to hypoxia, the main characteristic of CCRCC tumors [23]. Specifically, PAK2 was identified as the kinase with the highest kinase activation activity score (highest Z-score) that was unique to CCRCC. Survival analysis using The Human Protein Atlas of the kidney renal clear cell tissue confirmed our analysis of PAK2 as a unique key activating driver of CCRCC tumors. This demonstrates the capability of our model in identifying key tissue-specific drivers across tissues, lending it biological significance. With PAK2 confirmed having a central role in CCRCC, we sought to confirm it as a potential drug target. We successfully identified an FDA-approved SYK inhibitor, fostamatinib, that exhibits high binding affinity with PAK2 (Vinascore: −9.0 kcal/mol). Thus, this proves the extraordinary capabilities of SPARK in identifying new potential drug targets in the oncogenic signaling pathway.

### 3.3 Limitations and future work

We identified several areas that require attention in our future work:

**I. Class imbalance**: currently, class imbalance still exists in our ground-truth dataset, with a 1:5 positive to negative KSA ratio despite our efforts in mitigating them. This significantly impacts our AUPR results, lowering the effectiveness of our model (see Fig. 2). Therefore, it is required for us to increase the number of positive KSAs, such as using oversampling techniques or finding other reliable databases that can provide positive KSAs.
**II. Permutation test**: to further validate the results of SPARK beyond model performance scores, we cross-verified our prediction results with the Kinase Library, demonstrating excellent results. However, to determine the individual statistical significance of each edge and the overall statistical significance of each tissue-specific differential network, permutation testing would be required. However, due to limits in computational efficiency and time constraints, permutation testing was sadly not performed. In the future, permutation test would be required to validate our network on a more comprehensive scale.
**III. Tissue-specific differential networks**: as of now, we have not fully utilized both networks in junction to provide even more comprehensive analysis. Future work includes systematically integrating both networks to quantify edge-level and potential pathway-level differential activity, such as up- and down-regulation [14].

## 4 METHODS

### 4.1 Pan-cancer tissue-specific data

Data for SPARK was downloaded from the LinkedOmics Knowledge Base (LinkedOmicsKB, release 1): https://kb.linkedomics.org/download. LinkedOmicsKB is a pan-cancer, proteogenomics, data-driven knowledge base processed and precomputed from the Clinical Proteomic Tumor Analysis Consortium (CPTAC). We collected the proteomics and phosphoproteomics abundance matrices for Clear Cell Renal Cell Carcinoma (CCRCC), Colon Adenocarcinoma (COAD), Head and Neck Squamous Cell Carcinoma (HNSCC), Lung Squamous Cell Carcinoma (LSCC), Lung Adenocarcinoma (LUAD), Ovarian Cancer (OV), Pancreatic Ductal Adenocarcinoma (PDAC), and Uterine Corpus Endometrial Carcinoma (UCEC). Breast cancer (BRCA) and Glioblastoma (GBM) were excluded because the normal data was unavailable. The number of samples for each tissue are shown in Table 1, where the number of samples for each kinase and substate for each tissue is the same.

**Table 1:** Summary of the number of samples.

| # | Cancer type (tissue) | Number of normal samples | Number of tumor samples |
| --- | --- | --- | --- |
| 1 | CCRCC | 80 | 103 |
| 2 | COAD | 100 | 97 |
| 3 | HSNCC | 62 | 108 |
| 4 | LSCC | 99 | 108 |
| 5 | LUAD | 101 | 110 |
| 6 | OV | 19 | 83 |
| 7 | PDAC | 44 | 105 |
| 8 | UCEC | 18 | 95 |

### 4.2 Ground-truth dataset

We constructed the ground-truth dataset for SPARK by amalgamating experimentally verified results on kinase-substrate associations. Due to the scarcity of positive KSAs, the positive ground-truth dataset was formed by combining the positive training dataset for SELPHI 2.0, downloaded at: https://zenodo.org/records/13821321, and Phospho-SitePlus (PSP), downloaded at: https://www.phosphosite.org/staticDownloads.

To construct a set of negative samples, we first evaluated all possible kinase-substrate associations (KSAs) across all included cancer types based on the Kinase Library, available at: https://kinase-library.phosphosite.org/kinase-library/. KSAs with a kinase score percentile below 1 were initially selected as candidate negative instances. To mitigate potential bias introduced by specific phosphorylation sites that would cause class imbalance, we performed random sampling under the constraint that each site could appear at most five times in the negative dataset.

Due to variations in the number of quantified phosphorylation sites across different tissue types, the absolute numbers of both positive and negative samples differed among cancers. To address the severe class imbalance between positive and negative samples, we applied down-sampling to KSAs where the ratio of negative to positive samples exceeded 5:1. No further adjustment was made to the remaining associations.

### 4.3 Pairwise phosphoproteomic co-expression scores

To quantitatively characterize the potential linear dependencies between kinase-substrate interactions, we utilized pairwise phophoproteomic co-expression scores. For each cancer cohort, we first filtered out genes with less than 15 non-missing samples and per-gene missingness *≥* 40%. Median-imputation was applied on the filtered genes. For each kinase-substrate pair, we required a minimum of 15 overlapping samples to calculate the Pearson correlation coefficient (PCC) was calculated for each kinase-substrate pair in that cohort (Fig. 1). In addition, to ensure the reliability of the phosphorylation sites, we only retained the phosphorylation sites recorded in PhosphoSitePlus.

### 4.4 Kinase and phosphosite principal component representation

To calculate the principal component (PC) representation for each gene in each cohort, we first filtered out genes with pergene missingness *≥* 20%. Zero-imputation was used on the remaining genes and genes with zero variance were dropped. Single value decomposition (SVD) with Z-score scaled features is then applied to obtain PCs through the stats::prcomp function in R (Fig. 1).

### 4.5 STRING combined score

STRING protein network data for humans (version 12.0) was downloaded at: https://string-db.org/cgi/download. The combined scores were extracted and used (Fig. 1).

### 4.6 Phosphosite ESM-2 embeddings

The per-sequence embeddings were generated by compressing per-residue embeddings derived from the Evolutionary Scale Modeling 2 (ESM-2) protein language model. For each phosphosite, we extracted a 15 amino acid (AA) sequence (phosphosite residue flanked by 7 AAs) and passed them through esm2 t33 650M UR50D. esm2 t33 650M UR50D is a 33 layer, 650 million parameters ESM-2 protein language model. The 33rd (last) layer was then extracted and an autoencoder model was trained to compress the per-residue embeddings into per-protein embeddings to minimize data loss from conventional methods.

The autoencoder model compresses per-residue embeddings from a dimension of *n ×* 15 *×* 1280 to a dimension of *n ×* 1 *×* 20, where *n* represents the number of phosphosites. To achieve this, we constructed an encoder that compresses the embedding dimension from 1280 to 1, and a residue length reducer that compresses the 15 residues into 1 dimension. The latent layers (*n ×* 1 *×* 20) were extracted as features, providing a complex structural representation of the phosphosite and its neighbor. The autoencoder was trained on the mean-square error (MSE) of the results after expanding and decoding the latent layer. The ESM-2 embedding and autoencoder process can be visualized in Fig. 1b.

Normal and tumor cohorts were processed independently in ESM-2 and the autoencoder to produce separate embedding collections.

### 4.7 Hierarchical clustering of SPARK-derived network

To validate the capability of the SPARK model in identifying cancer-specific KSAs, we performed hierarchical clustering analysis on the inferred KSA networks. Specifically, a Euclidean distance matrix was computed based on the predicted KSA strengths across different cancer cohorts, followed by hierarchical clustering of samples based on this distance. For visualization purposes, 5000 kinase–substrate pairs were randomly selected to generate the heatmap, which was consistent with the clustering results obtained using the full dataset.

### 4.8 High priority kinase-substrate association filtering

KSAs with the top 1% of predicted probability and pair-wise kinase value *≥* 80 were filtered out as high priority KSAs.

### 4.9 Gene specificity metric (*τ* index) calculation

Tau (*τ*) index was used to quantitatively characterize gene specificity, calculated for both kinases and substrates in the tumor cohort [25]. For each high priority kinase or phosphosite, the normalized count (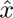) was calculated from the number of appearances (count: *x*) in the high priority group:

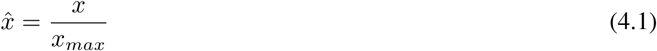

where *x_max_* = max(*x*_1_*, x*_2_*, x*_3_ *· · · x_n_*) and *n* is the number of tissues. *τ* was then calculated using 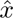:

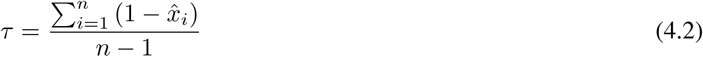

Thus, 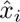 being 1 indicates less specificity and more conserved expression.

### 4.10 Weighted kinase-substrate enrichment analysis

Weighted Kinase-Substrate Enrichment Analysis (W-KSEA) was performed to evaluate kinase activity changes across cancer types. This approach integrates the confidence of kinase-substrate predictions as weights, providing a more robust estimation compared to conventional unweighted analyses.

For each cancer type, the global mean *µ_g_* and standard deviation *σ_g_* of the log_2_ fold-change (logFC) values for all predicted phosphosites were calculated. For a given kinase *k* with a set of *n* predicted substrates, the activity score was computed as a weighted Z-score. The weighted mean logFC (*µ_k_*) for the kinase was calculated as:

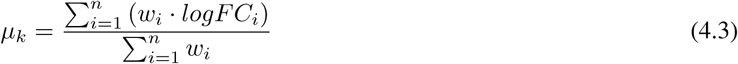

where *w_i_*is the predicted probability for substrate *i* being a true target of kinase *k* provided by SPARK. The effective substrate weighted count, *N_k_*, was defined as the sum of these weights:

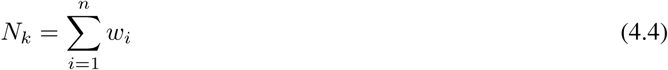

The kinase activity score (kinase-specific Z-score) was then derived by comparing the weighted mean to the global back-ground distribution:

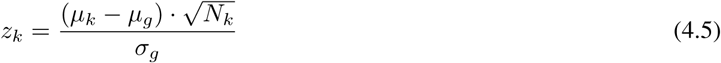

A two-tailed test was employed to calculate the p-value for each kinase. Kinases with a p-value of *<* 0.05 were considered statistically significant.

## 5 CODE AVAILABILITY

Code for SPARK is available at: https://github.com/AnsonZhang2009/SPARK_2025

## 6 DATA AVAILABILITY

Data used for the training of SPARK and produced by SPARK is available at: https://github.com/AnsonZhang2009/SPARK_2025

## ACKNOWLEDGMENTS

### Icons used in figures

The icons used in the figures were downloaded from Health Icons (https://healthicons.org/), Bioicons (bioicons.com), SMART medical art (smart.servier.com, and Scidraw (scidraw.io). All used icons are all free to use either with no credit required or under the CC BY-SA license.

### On the origin and development of SPARK

In 2018, the author’s whole family moved from Shenzhen to Guangzhou due to some devastating news - the seemingly normal stomach ache of the author’s grandfather was discovered to be stage 2 lung cancer (LSCC) caused by smoking. A year later, the desperation was made worse by the COVID-19 pandemic, which posed an even higher risk to my grandfather’s health, not only because it would couple with the existing lung cancer, but because it caused a lot of trouble in medical treatments. As a child back then, I was forced to watch my parents and grandmother care for my grandfather, the radiotherapy taking a toll on his physique.

10 years before my grandfather’s diagnostic of lung cancer, the author’s grandmother had caught liver cancer. However, due to determination and early diagnosis, she was able to successfully defeat it, though the author was too small to remember anything back then. Resection surgery and radiotherapy was performed on the author’s grandfather, and through sheer persistence and an optimistic attitude, he was able to defeat it.

Individual exploration and discovery led the author’s interests to be in the field of biochemistry and computer science, leading to the author’s attempt at finding biochemical-related potential solutions to mitigate the effects of cancer. Given the practical restrictions currently in place, such as the lack of laboratories to perform research, Dr. Shahid Ullah, one of my supervisors, suggested one specific field that amalgamated all three interests - using bioinformatics to analyze post-translational modifications (PTMs). Further analysis with Ms. Wendi Li, narrowed down the topic to kinase-substrate associations.

Through 8 months of arduous progress, we developed SPARK, a model for the systematic prediction of kinase-substrate relationships through statistical analysis and using online databases and R packages that are freely accessible. SPARK is the author’s first big attempt towards studying oncogenic processes, representing a tiny *spark* of hope, ignited by the author’s family history with cancer.

We would like to thank our two advisors: Dr. Shahid Ullah and Ms. Wendi Li from BASIS Bilingual School Shenzhen for providing the suggestions necessary for the construction of SPARK. Specifically, both helped in equipping the author with the necessary prerequisite knowledge in PTMs, statistical analysis, and writing refinement.

Lastly, we give our gratitude to the author’s parents, friends, and other teachers for supporting us through these 8 months of constant effort, and for renting the necessary server for me to perform statistical analysis. From the multiple failed times the server crashed on running ESM-2 to realizing that Y, T phosphosites were not included properly, it is they who have provided the support necessary for SPARK to continue.

